# Predicting nutrient profiles in food after processing

**DOI:** 10.1101/2022.09.28.509827

**Authors:** Tarini Naravane, Ilias Tagkopoulos

## Abstract

The future of personalized health relies on knowledge of dietary composition. The current analytical methods are impractical to scale up, and the computational methods are inadequate. We propose machine learning models to predict the nutritional profiles of cooked foods given the raw food composition and cooking method, for a variety of plant and animal-based foods. Our models (trained on USDA’s SR dataset) were on average 31% better than baselines, based on RMSE metric, and particularly good for leafy green vegetables and various cuts of beef. We also identified and remedied a bias in the data caused by representation of composition per 100grams. The scaling methods are based on a process-invariant nutrient, and the scaled data improves prediction performance. Finally, we advocate for an integrated approach of data analysis and modeling when generating future composition data to make the task more efficient, less costly and apply for development of reliable models.

## INTRODUCTION

Food processing, such as fermentation, baking, or even boiling alters the chemical composition of food, often in unpredictable ways from the raw to the finished state. This is due to the unresolved chemical and structural complexity of the food and the physio-chemical transformation mechanisms that occur during processing. [1] [2] Yet in spite of these challenges, the objectives for prediction models are compelling which include sensory properties [3] such as aroma, texture, taste, etc., and nutrient profiles, and here we address the latter. The models in use currently, simplify the inherent complexity and instead predict the content of a nutrient based on only a few parameters. For instance, kinetic modelling based on experimental data for any given food establishes the relationship between nutrient concentration, time and temperature conditions. [4] [5] [6] [7]. This can then be applied to compute concentrations, for example predicting vitamin C (ascorbic acid) content in processed orange juice [6]. Another approach to compute post-process nutrition composition, is to apply retention factors (RF) which are based on analytical composition data on a representative set of foods and processes. RF-based computation is used widely by food manufacturers for nutrition labels, and by USDA’s dietary survey group to calculate nutrient intakes that investigators may use to determine correlations between intake and health outcomes [8]. However, all of these methods have limited potential. Kinetic models are difficult to scale up to capturing more food and processing parameters, as these measurements are time-consuming, expensive [9] and have many experimental challenges such as certain chemicals which degrade rapidly. RF-based methods in practice inevitably under or overestimate the nutrient content in a particular instance, since any single RF is representative of several foods and a cooking method. Here we address the challenge that our knowledge of composition and reactions of food systems is limited, which inevitably manifests to such incomplete or underdetermined models. This can be at least partially addressed with predictive machine learning (ML) methods that can learn the multi-parametric transformation patterns between the compositions of raw and cooked foods, from experimental data across diverse foods and cooking methods.

The application of ML to food science data is at an early stage, yet it has been successful in generalizing across a variety of prediction tasks when trained on relevant datasets. Some examples of recent work include prediction of nutrient profiles or properties of food. Classifiers model have been applied to predict sensory properties from the molecular structure, such as bitter [10] [11] and sweet [12] [13]and aroma labels [14]. A number of food quality classifiers use hyperspectral data, for example for the freshness classification of shrimp [15], detection of adulteration in red meat products [16] or detection of damaged/bruised fruits and vegetables [17] [18]. Several models have addressed attributes related to nutrient profiles. Natural language processing (NLP) methods were used to predict the macronutrient (proteins, fats and carbohydrates) content of foods from a text description of the food [19]. USDA investigators predicted the content of 3 label nutrients (carbohydrates, protein and sodium) in processed foods from the ingredient list, using the Branded Foods datatype in Food Data Central (FDC) [20]. Several projects predicted nutrient contents from the composition data; nutrient content was predicted for the missing values in food composition data [21], lactose content was predicted in dietary recall database [22], fiber content was predicted for commercially processed foods [23]. Availability of datasets with high quality data for training and testing is essential, and databases such as BitterDB [24], BTP640 [25], FlavorDB [26],FooDB [27], SuperSweet [28], Fenaroli [29],GoodScents [30], FDC [31] as well as specifically curated datasets of hyperspectral images may contribute to this end.

Here, we have constructed an ML model that predicts food micronutrient (specifically 7 vitamins and 7 minerals) composition after processing (**Figure** 1). We have curated a sample of 820 foods, for 5 processes, namely steaming, boiling, grilling, broiling, and roasting from FDC, and trained regressors per nutrient and per process that have achieved a correlation(R^2^) between measured and predicted micronutrient values that range from 0.42 to 0.95 (outliers are -0.42, -0.09,0.13 and 0.23).

**Figure 1.**
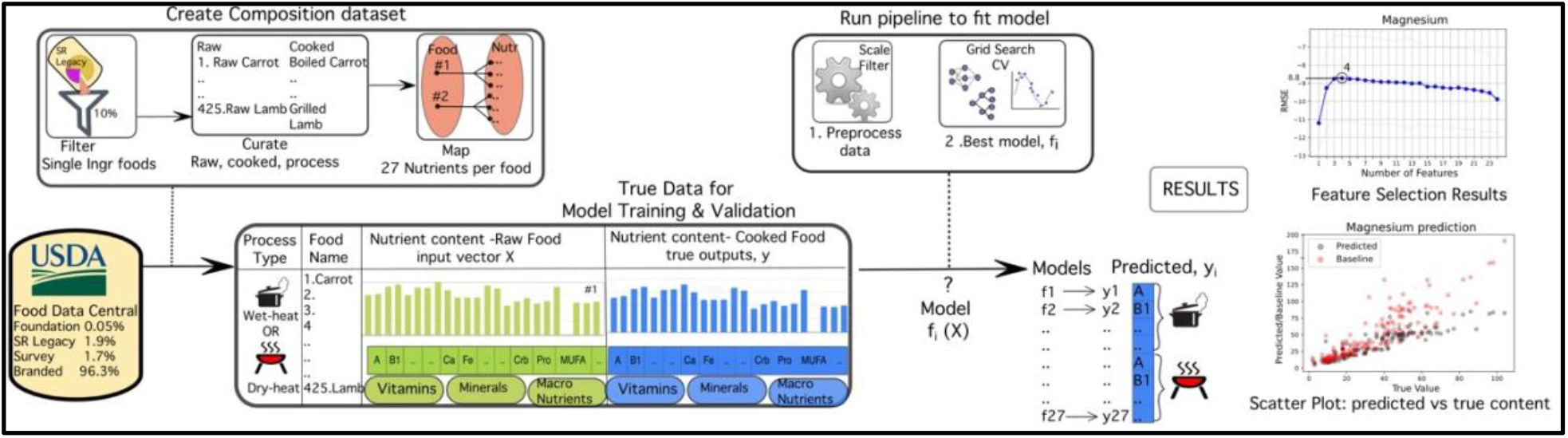
Overview of architecture (left to right) from data selection to prediction results. Single ingredient foods are selected from SR legacy (one of the 5 data types in FDC), and then organized by pair (raw,cooked) and process type. Cooking processes include boiling and steaming which are grouped into wet heat processes(WH) and broiling, grilling, and roasting which are grouped into dry heat processes (DH). Foods are mapped to composition, with 27 components per food. Models are trained from composition data, such that the input feature is the composition of the raw food, and each model is trained separately for every micronutrient in the cooked food. Models are trained separately for both process types, with 14 for WH and 13 for DH (excluding vitamin C predictor model). Prior to model fitting, the composition data is scaled and filtered. Model fitting uses a grid search cross validation approach, such that there are 12336 regressor models. The best model has the least error, RMSE. Then predicted composition is compared to the actual (ground truth) composition in two results. The scatter plot for prediction of magnesium content shows the both the prediction (black dots) and baseline (red dots) values on the Y axis, versus the actual values (X axis). The other result is the performance (RMSE) analysis against the feature (input features) size.

## METHODS

### Dataset

We downloaded the composition dataset of 7,793 foods from the Standard Reference (SR) legacy dataset, which is the most suitable of the 5 data types in FDC (**Figure 1**; as of November 2021), since it is aligned with our objectives. This dataset is intended for application in public health initiatives such as the assessment of nutrient intakes for the purpose of national nutrition monitoring, in creating meal plans in schools and day-care centers, in product development and labeling by manufacturers. The composition data for the foods in SR is obtained from 3 sources; analytical experiments, calculations (based on the analytical data), and literature. For our models, we selected a subset of these data according to the following criteria. We matched raw/cooked food pairs, where the raw foods were a single ingredient harvested from a plant or from an animal (includes butchery products), and the cooked food was the outcome of the raw food treated to wet (boiling, steaming), or dry (roasting, grilling, broiling) heat processes. Foods were excluded from the dataset if either there was no single-ingredient raw food corresponding to the cooked food and vice-versa, or the foods had several ingredients and produced by a multi-step process like ‘Luncheon meat, pork and chicken, minced, canned, includes SPAM Lite’, ‘Bread, banana, prepared from recipe, made with margarine’. We excluded processes which have added ingredients, such as oil for frying, while we included boiling and steaming (simple aqueous, i.e., wet heat processes), as well as roasting, broiling and grilling (dry heat processes). This resulted in 840 foods total in the dataset, with 178 and 247 pairs from wet and dry heat processes, respectively. In this dataset, all plant-based foods were cooked by wet heat process (WH), and all animal-based foods by a dry heat (DH) process. The categorical breakdown of the number of pairs for plant-based and animal-based foods is shown in **Figure 2**.

**Figure 2.**
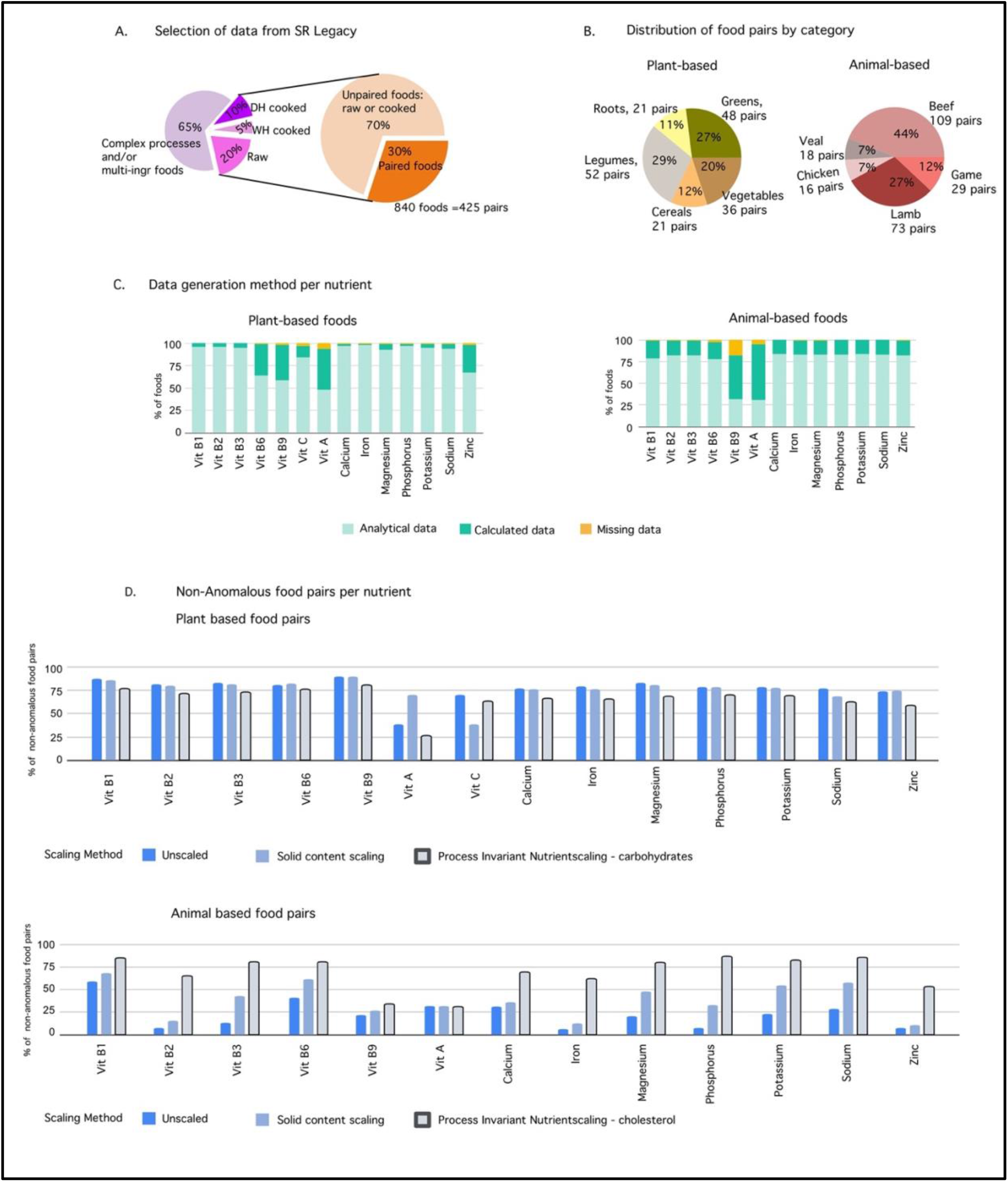
Data Review. (A) Out of 7,793 foods in the SR Legacy datatype in FDC dataset, 2,724 (35%) are single ingredient foods. Within that set, we identified 425 pairs of raw-cooked single ingredient foods. (B) The food pairs per category for plant-based and animal-based foods. There are a total of 178 pairs of plant-based foods and 247 pairs of animal-based foods. (C) The food-pair distribution by the method of data generation. (D) Comparing the percentage of food-pairs of non-anomalous data by scaling method.

The composition data consists of content values for up to 232 ‘chemical constituents’ or ‘components’, which include specific chemicals (vitamins, amino-acids, fatty acids, etc.) and aggregated chemicals or chemical groups (total fats, total proteins, etc.) for every food. Here, we selected the components that are reported for at least 80% of the foods in our dataset. This resulted in 27 components per food, namely 9 vitamins, 10 minerals, water, and 7 aggregates of total protein, total carbohydrates and various fat categories (**Supplementary materials**). The pair-wise composition data was used to train the prediction models where the input feature set to every model is the content of the 27 components in the raw food and the outputs are the contents of the 14 micronutrients in the cooked food. Prior to model fitting, the composition data should be preprocessed to adjust for the bias resulting from the conventional format of representing nutrient contents per 100 grams of a food sample. In practice, the raw sample is likely not always 100 grams, and the cooked food sample would have a relatively higher or lower weight yield primarily due to loss or gain of water in the cooking process. As a result, when water is lost the solid components have a relatively higher concentration, and in comparing the raw and cooked food composition per 100 grams, there is a concentration bias in the cooked food. Similarly, there is a dilution bias when water is absorbed in the cooking process. Ideally the data preprocessing would reverse this scaling effect. We use two different scaling methods, solid content scaling in equations 1 and 2 and process-invariant nutrient scaling in equations 3 and 4. For the solid content scaling (SCS), the water content in the raw food is set to that in the cooked food which assumes that the water content stays constant while scaling the content of other components. Then the composition is scaled preserving the proportions and the weight of the sample is maintained at 100 grams. This method lowers the extent of the dilution or concentration biases while the “process-invariant nutrient” scaling (PINS) method intends to undo this bias. The method is based on identifying a nutrient that is largely invariant to processing and can be used to calculate the scaling factor as per equation 3. This factor is then used as per equation 4 to derive the composition for the “true” weight of the cooked food corresponding to a 100-gram sample of raw food. From the 27 components in our dataset, cholesterol, iron and zinc are invariant to changes in processing for animal-based foods, but there is no such information for the plant-based foods. Cholesterol in the various meats is invariant to processing since it is in the muscle-cell membranes which are resistant to cooking loss. [32]. Iron and zinc were reported to be process invariant as per experimental studies conducted by UDSA [33]. For confirmation of these hypotheses, all components are used in the PINS method and prediction performance is compared for both animal and plant-based foods.

In the equations for scaling methods, R represents the raw food and C represents the cooked food, and component is the generalized term for each of the 26 solid components.

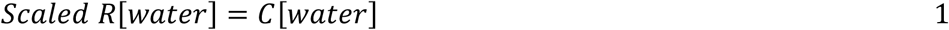

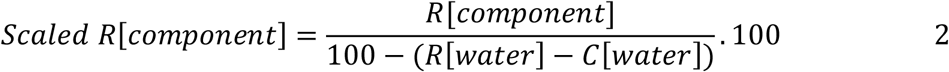

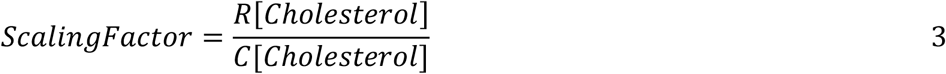

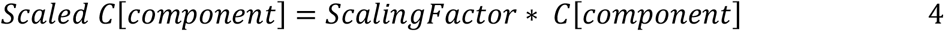

The final dataset includes 425 pairs of foods, with 27 components, 5 processes (boiling, steaming, roasting, grilling and broiling), in two states (raw and cooked). **(Supplementary materials)**.

## MODEL

We trained models to predict the content of 14 micronutrients for which we had baseline retention factors in the cooked food. Of those, seven are vitamins, namely vitamin B1 (thiamin), vitamin B2 (riboflavin), vitamin B3 (niacin), vitamin B6 (pyridoxine), vitamin B9 (folate), vitamin C (ascorbic acid), vitamin A, and the other seven are minerals, namely calcium, iron, potassium, phosphorus, magnesium, sodium and zinc. We created separate models based on the process category (wet, dry), as these are fundamentally different processes, but not based on the actual process (e.g., boiling vs. steaming), as there are not sufficient data per process to avoid overfitting. All models have the same input, which is the composition of the raw food, as illustrated in **Figure 1**. Other details that might be informative to the task (cooking time, temperature, water content) were not available in the SR legacy dataset, and consequently were not present in our dataset, or our model. Since vitamin C is not present in meats (which are all the foods for DH models), the dry heat models are only 13, for the other micronutrients, resulting in 27 models total (13 for DH and 14 for DH). These sets of WH and DH models were trained and tested on scaled variants of the dataset explained earlier. We applied a filtering step to the scaled datasets to select the pairs of foods where the nutrient being predicted was more in the raw food than in the cooked food. So, each of the nutrient models were trained on different subsets of the data. The unscaled data for the dry heat models and wet heat models was not filtered for this condition. The effect of the data scaling and filtering on the predictive models is explained in the **Results**.

The best performing model (for any dataset variant) was selected based on a cross validation grid search across 6 regressor types (MLP, LASSO, Elastic Net, Gradient Boost, Random Forest, Decision Trees), each with a variety of hyperparameters totaling 12,336 regressors where the metric for the best model was the least root mean squared error (RMSE). This was done for each of the 27 models using the sklearn library [34] and the best hyperparameters for each of the regressor types along with the RMSE is in **Supplementary materials**. We then performed a feature selection technique, a recursive feature elimination variant as described in the sequential feature selector function of the mlxtend package [35]. The model performances for data variants for the WH and DH process are compared in **Table 1**.

**Table 1.** Comparing models trained on different data variants. The prediction performance results for the models trained on data variants specified in the Methods are shown in this table. The metric for model performances is RMSE – Root mean squared error. A complete coverage of all performance for all PINS data is in **Supplementary materials**. Data Variants: Unscaled is the original data. SCS – Solid content scaling. PINS – Process Invariant Nutrient scaling and the specific nutrients is in parenthesis.

| OUTPUT | WETHEAT |  | DRYHEAT |  |  |  |  |
| --- | --- | --- | --- | --- | --- | --- | --- |
|  | Unscaled | SCS | Unscaled | SCS | PINS ( Zinc) | PINS(Iron) | PINS(Cholesterol) |
| Thiamine | 0.04 | 0.03 | 0.03 | 0.02 | 0.01 | 0.01 | 0.01 |
| Riboflavin | 0.05 | 0.04 | 0.05 | 0.03 | 0.02 | 0.02 | 0.02 |
| Niacin | 0.48 | 0.25 | 0.87 | 0.38 | 0.46 | 0.60 | 0.45 |
| B6 | 0.05 | 0.04 | 0.09 | 0.04 | 0.07 | 0.05 | 0.06 |
| Folate | 22.37 | 20.53 | 4.38 | 8.39 | 6.46 | 3.74 | 1.72 |
| VitC | 13.28 | 8.28 | NA |  |  |  |  |
| VitA | 83.21 | 23.85 | 3.37 | 1.7 | 1.25 | 2.81 | 1.5 |
| Calcium | 22.17 | 25.76 | 4.33 | 3.4 | 2.41 | 1.36 | 1.81 |
| Iron | 0.6 | 0.45 | 0.33 | 0.3 | 0.14 | 0.00 | 0.16 |
| Magnesium | 11.19 | 8.83 | 5.05 | 2.4 | 2.19 | 2.22 | 2.33 |
| Phosphorus | 24.1 | 22.19 | 24.16 | 21.8 | 15.60 | 15.67 | 21.41 |
| Potassium | 101.9 | 72.13 | 48.87 | 42.3 | 27.49 | 30.95 | 32.23 |
| Sodium | 17.4 | 13.44 | 13.26 | 8.7 | 6.80 | 7.35 | 9.36 |
| Zinc | 0.2 | 0.2 | 0.67 | 0.3 | 0.00 | 0.38 | 0.29 |
| AVERAGE | 21.22 | 14 | 8.11 | 6.9 | 5.24 | 5.43 | 5.49 |

We assessed the predictive performance (RMSE) in comparison with 2 baseline models. The first is to naively assume that the dependent variable (the micronutrient to predict after cooking) is equal to its value in the raw food. This baseline serves as a comparison to a naïve regressor where the retention factor (RF) is 100%, i.e., the amount of the micronutrient after the heat process is the same as in the raw food. The second baseline was based on the USDA Retention Factors table, a common, standard model for the retention of nutrients after a process [8]. The nutrient outputs were computed as a product of the RF for the specific nutrient and the content of that nutrient in the raw food. We use RSME, the coefficient of determination (R^2^), Pearson Correlation Coefficient (PCC), and Spearman Rank Correlation Coefficient (SRC) to assess the performance of our regressor model (**Table 2** and **Supplementary materials**). At each case, we performed 5-fold cross validation runs, bootstrapped 50 times to avoid overfitting and increase the generalization potential of our classifiers. For a subset of foods (**Supplementary materials**), we provide a higher resolution baseline using retention factors from experimental studies in literature. Finally, we analyze the prediction performance through a breakdown of R^2^ by food category for plant-based foods (Leafy greens, Roots, Vegetables, Legumes, Cereals) and animal-based foods (Beef, Lamb, Chicken, Veal) as shown in **Table 3**. We do this by tagging every predicted micronutrient value by the category (associated with the food) and calculate the R^2^ for every group. This is repeated for all predictions, and the average R^2^ of a category is used to determine the best and worst performances in the plant-based and animal-based foods.

**Table 2.**
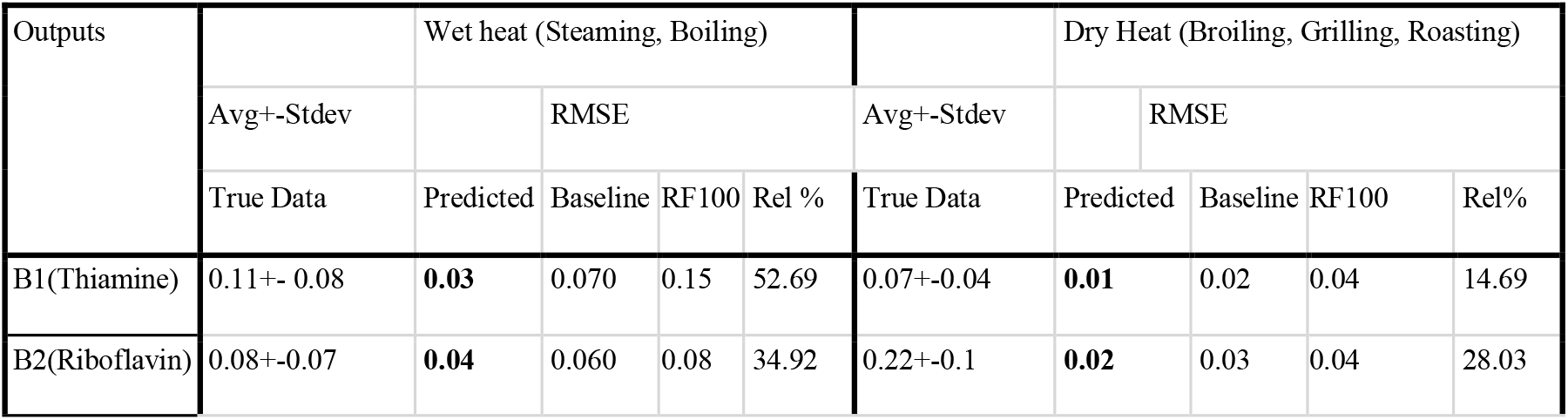
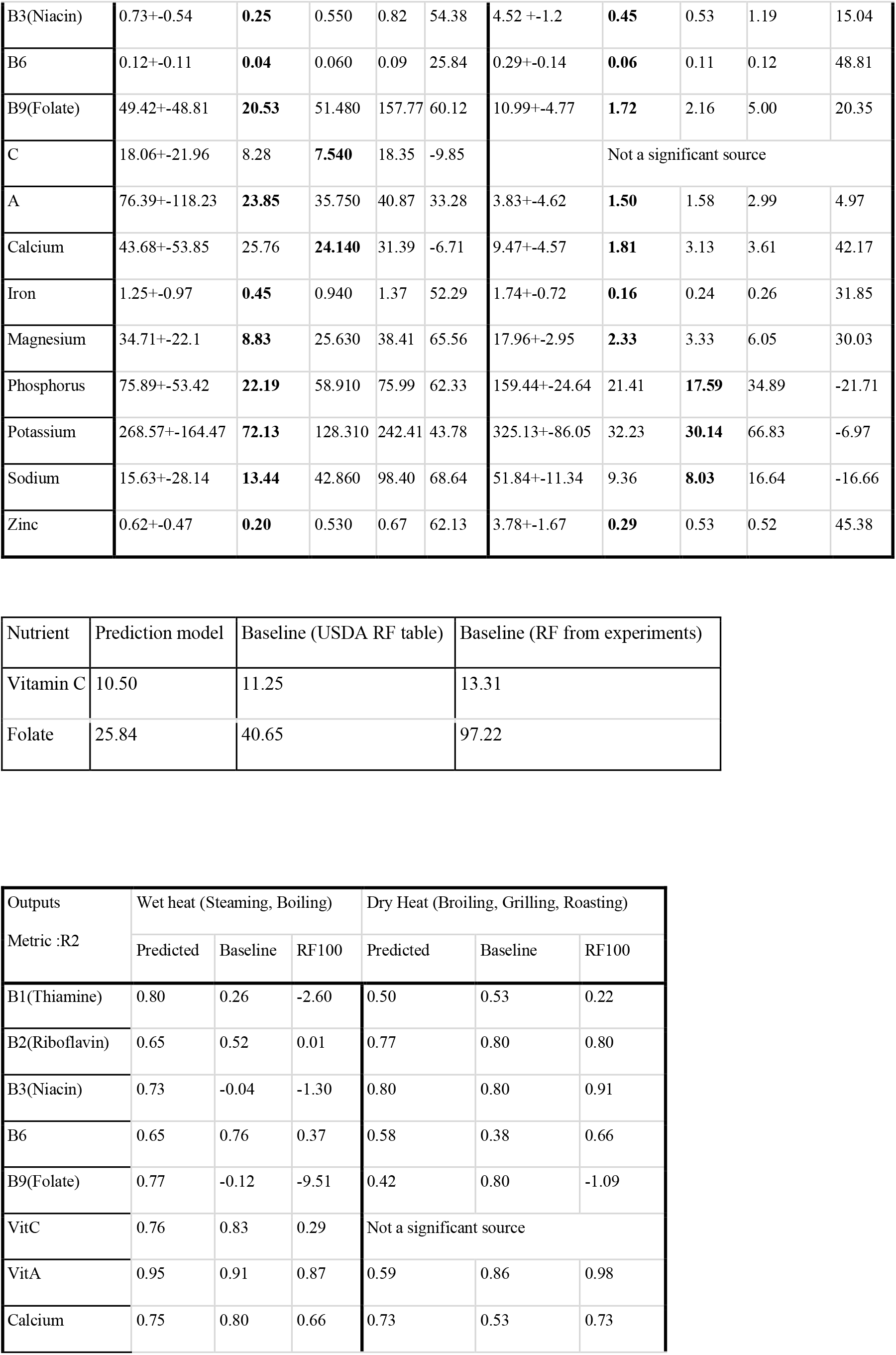
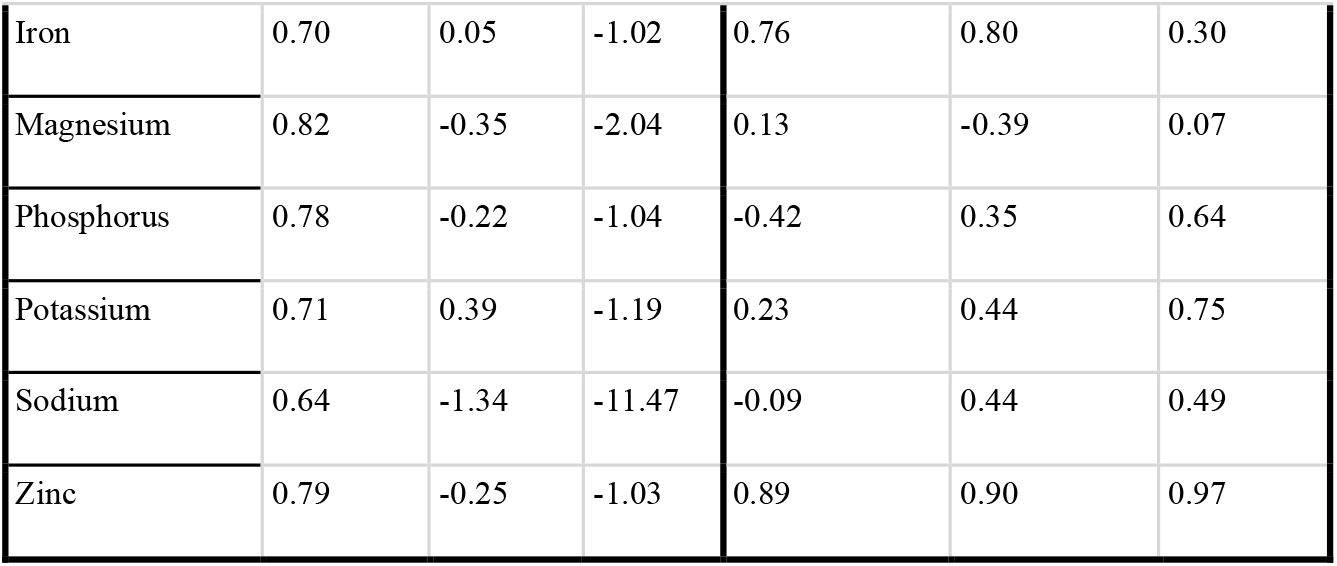
Results of prediction models compared to baselines. The prediction scores (RMSE and R^2^) are the average of 50 runs, due to the inherent randomness in the models. [A] (RMSE) of best prediction models, compared to baseline (USDA’s RF guide Version 6) model and naïve model (output content=input content). The better of the prediction or baseline score is highlighted. The rel% column is calculated as : (baseline-predicted)/baseline*100 1B. Additional baseline model for vitamin C (ascorbic acid) and vitamin B9 (folate) using RF values from experiments on selected foods. 1C. The metric R^2^ (coefficient of determination) is scale invariant (as opposed to the RMSE) for ease in comparison across all predictions. The corresponding box plot is in Figure 3.

**Table 3.** Various metrics (R^2^, RMSE,PCC) by category for plant-based foods. Cereals do not have data for vitamin A and C predictions. Abbreviations are used for the predicted nutrient, Ca:Calcium, Fe:Iron, Mg:Magnesium, Ph:Phosphorus, K:Potassium, Na:Sodium, Zn:Zinc. The remainder are vitamins.

| Output | Leafy Greens |  |  | Roots |  |  | Vegetables |  |  | Legumes |  |  | Cereals |  |  |
| --- | --- | --- | --- | --- | --- | --- | --- | --- | --- | --- | --- | --- | --- | --- | --- |
| | RMSE | $R^2$ | PCC | RMSE | $R^2$ | PCC | RMSE | $R^2$ | PCC | RMSE | $R^2$ | PCC | RMSE | $R^2$ | PCC |
| Ca | 23.97 | 0.82 | 0.90 | 7.54 | 0.90 | 0.95 | 73.06 | 0.41 | 0.98 | 21.88 | 0.55 | 0.88 | 7.57 | 0.50 | 0.92 |
| Fe | 0.29 | 0.82 | 0.90 | 0.24 | 0.56 | 0.75 | 0.15 | 0.76 | 0.86 | 0.69 | 0.49 | 0.71 | 0.39 | 0.52 | 0.77 |
| Mg | 8.95 | 0.80 | 0.89 | 4.65 | 0.83 | 0.96 | 5.12 | 0.85 | 0.93 | 10.72 | 0.70 | 0.84 | 11.03 | 0.58 | 0.81 |
| Ph | 11.37 | 0.66 | 0.83 | 10.33 | 0.82 | 0.89 | 24.90 | 0.44 | 0.58 | 26.92 | 0.65 | 0.85 | 27.41 | 0.48 | 0.79 |
| K | 76.50 | 0.70 | 0.84 | 71.64 | 0.88 | 0.94 | 68.81 | -0.12 | 0.72 | 73.35 | 0.49 | 0.79 | 58.98 | -0.19 | 0.27 |
| Na | 23.85 | 0.68 | 0.80 | 9.90 | 0.81 | 0.91 | 4.92 | 0.50 | 0.98 | 9.02 | 0.11 | 0.88 | 19.06 | 0.78 | 0.92 |
| Zn | 0.10 | 0.72 | 0.86 | 0.09 | -0.23 | 0.59 | 0.08 | 0.96 | 0.97 | 0.32 | 0.48 | 0.70 | 0.23 | 0.68 | 0.83 |
| A | 36.24 | 0.93 | 0.97 | 19.73 | 0.65 | 0.96 | 21.48 | 0.91 | 0.98 | 11.66 | 0.45 | 0.76 | NA | NA | NA |
| C | 11.97 | 0.65 | 0.84 | 6.65 | 0.85 | 0.94 | 7.05 | 0.88 | 0.96 | 3.71 | 0.77 | 0.94 | NA | NA | NA |
| B1 | 0.02 | 0.84 | 0.92 | 0.01 | 0.92 | 0.99 | 0.01 | 0.85 | 0.92 | 0.05 | 0.64 | 0.80 | 0.04 | 0.66 | 0.85 |
| B2 | 0.06 | 0.64 | 0.81 | 0.04 | 0.28 | 0.67 | 0.03 | 0.76 | 0.88 | 0.02 | 0.65 | 0.85 | 0.04 | 0.61 | 0.79 |
| B3 | 0.16 | 0.76 | 0.93 | 0.12 | 0.82 | 0.92 | 0.27 | 0.69 | 0.82 | 0.19 | 0.64 | 0.85 | 0.49 | 0.69 | 0.85 |
| B6 | 0.06 | 0.90 | 0.95 | 0.03 | 0.79 | 0.95 | 0.04 | 0.63 | 0.86 | 0.04 | 0.56 | 0.70 | 0.03 | 0.54 | 0.79 |
| B9 | 20.90 | 0.63 | 0.80 | 7.18 | 0.82 | 0.97 | 4.90 | 0.89 | 0.95 | 29.38 | 0.68 | 0.83 | 19.88 | 0.43 | 0.91 |

**Various metrics ( $R^2$ , RMSE, PCC) by category for animal-based foods**
| Output | Beef |  |  | Lamb |  |  | Chicken |  |  | Veal |  |  |
| --- | --- | --- | --- | --- | --- | --- | --- | --- | --- | --- | --- | --- |
| | RMSE | $R^2$ | PCC | RMSE | $R^2$ | PCC | RMSE | $R^2$ | PCC | RMSE | $R^2$ | PCC |
| Ca | 1.85 | 0.76 | 0.88 | 1.93 | 0.87 | 0.83 | 2.27 | 0.04 | 0.59 | 3.52 | 0.68 | 0.78 |
| Fe | 0.13 | 0.89 | 0.95 | 0.16 | 0.25 | 0.67 | 0.08 | 0.84 | 0.95 | 0.22 | -1.63 | 0.34 |
| Mg | 2.68 | 0.03 | 0.48 | 1.50 | 0.31 | 0.57 | 2.22 | 0.43 | 0.67 | 4.48 | 0.10 | 0.37 |
| Ph | 18.31 | 0.32 | 0.70 | 19.91 | -0.44 | 0.48 | 25.51 | -0.06 | 0.73 | 46.28 | -1.86 | -0.12 |
| K | 33.23 | 0.19 | 0.60 | 32.69 | 0.31 | 0.63 | 32.81 | 0.15 | 0.88 | 37.47 | 0.25 | 0.91 |
| Na | 9.50 | 0.25 | 0.65 | 7.15 | -0.27 | 0.35 | 13.65 | -0.51 | 0.46 | 14.97 | -1.00 | 0.33 |
| Zn | 0.38 | 0.95 | 0.98 | 0.24 | 0.91 | 0.96 | 0.15 | 0.80 | 1.00 | 0.17 | 0.93 | 0.99 |
| A | 1.24 | 0.39 | 0.90 | 1.67 | 0.80 | 0.91 | 3.67 | 0.73 | 0.86 |  |  |  |
| B1 | 0.01 | 0.60 | 0.80 | 0.01 | 0.73 | 0.87 | 0.01 | 0.62 | 0.92 | 0.02 | -0.29 | 0.56 |
| B2 | 0.02 | 0.95 | 0.98 | 0.01 | 0.94 | 0.97 | 0.03 | -0.10 | 0.94 | 0.06 | -0.06 | 0.47 |
| B3 | 0.47 | 0.79 | 0.89 | 0.40 | 0.70 | 0.84 | 0.85 | 0.82 | 0.95 | 0.41 | 0.92 | 0.97 |
| B6 | 0.05 | 0.78 | 0.88 | 0.04 | 0.89 | 0.95 | 0.02 | 0.83 | 0.95 | 0.15 | -1.24 | 0.47 |
| B9 | 0.99 | -0.63 | 0.54 | 1.74 | 0.67 | 0.83 | 0.91 | -2.81 | -0.64 | 2.62 | -2.82 | 0.91 |

## RESULTS

### Approximately 10% of SR Legacy foods can be paired in processes to be used in model training

The single ingredient foods that are either raw or cooked were found in 35% of the SR legacy data, and 30% of these were paired into raw and cooked samples. The final selection of 840 foods (or 425 pairs) is 10% of SR legacy data (**Figure 2A**), with an unequal distribution of data pairs by food category (**Figure 2B**). We identified an anomaly where the content of a micronutrient was more in the cooked food than in the raw food in 50% of the pairs on average across the 14 micronutrients. The anomaly was more severe for the animal-based foods (77% vs 23% pairs, respectively; see **Supplementary materials**). This was partially caused by the bias introduced by the data representation convention. For the animal-based foods, the non-anomalous pairs are 30% of the total pairs for unscaled data and increase to 70% for PINS-cholesterol scaled data, p-value<0.01. This is reasonable, since the anomaly is due to a concentration bias (nutrient content in cooked food is more than in raw food), which is mitigated by scaling. For the plant-based foods, there is no significant change(p-value>0.05) in non-anomalous pairs using the scaling methods for plant-based foods, since the issue is a dilution bias which is mitigated however this does not cause an anomaly (nutrient content in cooked food is more than in raw food). The comparison of non-anomalous pairs for animal and plant-based foods is shown in **Figure 2**. The **Discussion** section explains the reasons for this differing effectiveness of the scaling methods in reducing the bias and suggests other possible causes for the bias.

### Scaling improves model performance

We trained predictive models on variants of the datasets as explained in Methods. The dry heat models (broiling, grilling, roasting processes; 247 animal-based foods) and wet heat models (steaming, boiling; 178 plant-based foods) were trained on the unscaled data, which is not filtered for the anomalous condition, and on data from both the scaling methods which is filtered for non-anomalous data. We use the metric RMSE to compare model performance and confirm the hypotheses described in Methods. For the dry heat models, the average RMSE (for 13 predictions) was 20% lower when the model was trained on data scaled by the PINS-cholesterol method than data scaled using SCS method, which had 15% lower RMSE compared to the model trained on unscaled data. Although the model performance based on PINS data for iron and zinc has lower average RMSE than cholesterol, we consider the model trained on PINS-cholesterol as the best model since there is a mechanistic explanation described in **Methods**. For the wet heat models, the average RMSE was 35% lower when the model was trained on SCS data than that on unscaled data. These comparisons are shown in **Table 1**, and all results are in **Supplementary materials** and further analysis is in **Discussion**. The best model for the wet heat process is trained on SCS data and for the dry heat process it is trained on PINS-cholesterol data. We now compare results from the best predictive ML models to the baseline model.

### The predictive model performs 43% and 18% better than using the standard USDA Retention Factor model for wet and dry heat processes, respectively

We compared the predicted concentrations of the micronutrients in the cooked foods for both the wet heat processes and the dry heat processes against the two baseline models, as described in the **Methods** section. When compared to the naïve baseline (i.e., retention factor is always 100%), the predictive model is better in all 27 out of the 27 comparisons (100%; RMSE of 9.90±16.45 vs 31.29±56.56, respectively; 64% decrease of RMSE on average for wet heat, p-value < 0.01; 52% decrease in RMSE on average for dry heat, p-value < 0.01). Then, to compare with the standard practice, we computed micronutrient concentrations using the USDA’s Retention Factor table (see **Methods**) as shown in **Table 2**. In that case, the predictive model was better than this baseline in 22 out of the 27 comparisons (81%; RMSE of 9.90±16.45 vs 16.45±28.41, respectively; 43% decrease of RMSE on average for wet heat, p-value < 0.01; 18% decrease in RMSE on average for dry heat, p-value < 0.01). **Figure 3** depicts the correlation between predicted and actual (ground truth) values for the 14 micronutrients, for both the ML model and the USDA retention factor baseline. Next, we investigated the difference in the predictive performance when curating retention factors from literature. For this, we identified the retention factors of vitamin C (ascorbic acid) for 12 sample foods (green beans, beet greens, broccoli, Chinese cabbage, carrots, cauliflower, mustard greens, green peas, green peppers, pumpkin, spinach, zucchini) and of vitamin B9 (folate) for 12 sample foods (amaranth leaves, broccoli, drumstick leaves, snap beans, lentils, okra, onions, potatoes, green peas, soybeans spinach, taro leaves) (see **Supplementary materials**). In both cases, the ML model had a better agreement with the ground truth data than the Literature Retention factor baseline, although less so for vitamin C (for vitamin C (ascorbic acid), RMSE 10.51 vs 13.31, p-value=0.026; for vitamin B9 (folate) RMSE 25.84 vs 97.22, p-value=0.013). Note that retention factor information for each micronutrient is not available for the majority of foods, and it is a time consuming and expensive process to measure it. Using scale-invariant metrics reach the same conclusions (see **Supplementary materials)**. The **Discussion** section elaborates further on the reasons that any RF baseline method is error prone and not appropriate to compute nutritional baselines.

**Figure 3.**
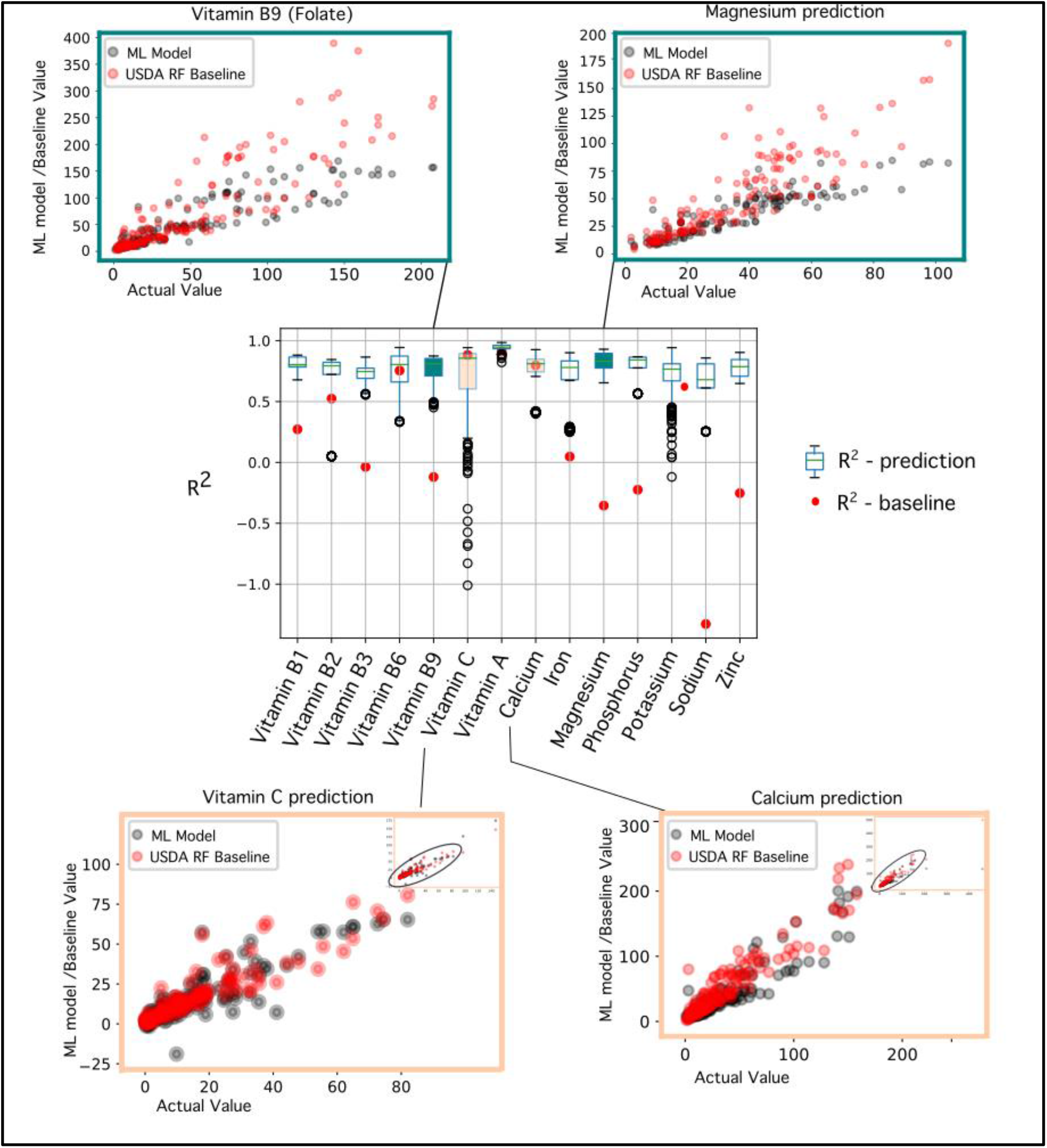
Model Performance Analysis. Centre: Box plot of R^2^ (coefficient of determination) for 14 prediction models and corresponding USDA baseline model. Scatter plots for values from the prediction models and USDA baseline model, against actual values (ground truth) are shown in both cases where prediction was better or worse than the baseline. The top 2 scatter plots are for the case where prediction was better. Plots for vitamin B9 (folate) and magnesium show that the baseline model tends to have erroneously higher values than the predicted values, relative to the actual data. Lower 2 scatter plots are for the case where prediction was worse than baseline by only a small margin. Plots for vitamin C and calcium have a noticeable overlap in values for the prediction model and baseline.

### Prediction performance is best for leafy green vegetables, and worst for cereals, in the plant-based food categories, and best for beef and worst for veal in the animal-based food categories

As reported in prior literature, the food structure/phenotype influences the chemical and physical changes that occur in food processes. Here we use the food category to represent this concept and show the differences in predictability. We group the 14 predicted micronutrient values by the food category and calculated the R^2^ (**Table 3** and **Figure 4A**). Leafy green vegetables have the highest average R^2^ of 0.75±0.10 and cereals have the least average R^2^ of 0.52±0.25. In the dry-heat processed animal-based foods, beef had the highest average with R^2^ of 0.48±0.46 and veal the least average R^2^ of - 0.50 ±1.21. Due to the uncertainty associated with methods of data generation, USDA specifies the nutrients with most reliable data, these are vitamin B3 (niacin), vitamin B6, calcium, iron and zinc. The highest average R^2^ is now 0.85 ±0.08 for beef and the lowest is -0.06 ±1.26 for veal. As such, the nutrient loss is better predicted in leafy greens and beef given the current training data.

**Figure 4:**
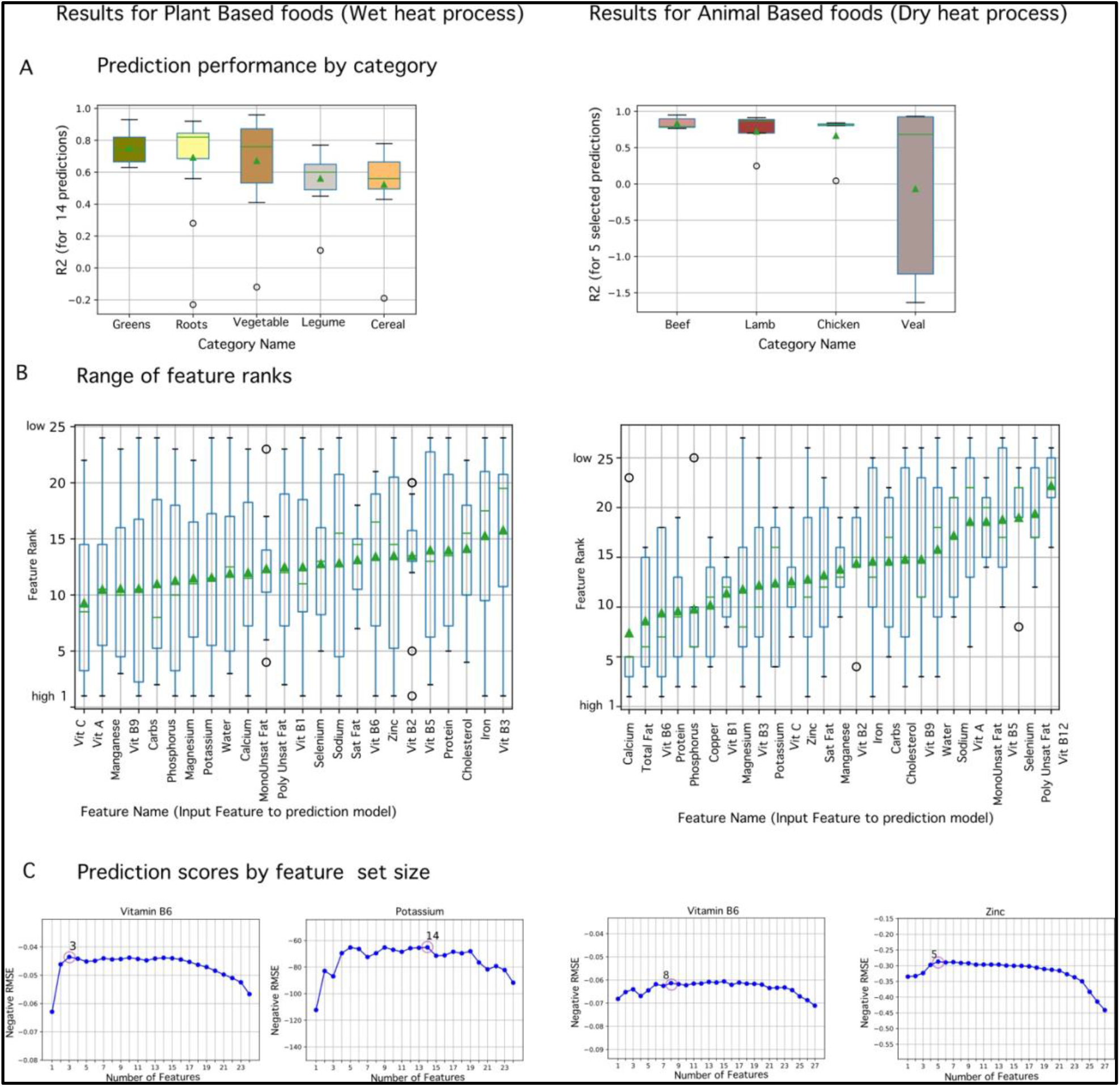
Results. (A): Box plot of R^2^ for predictions by food category. For the plant-based foods the box plot shows all 14 predictions. Leafy green vegetables have the best performance and Cereals the worst. For the animal-based foods, only 5 predictions are considered since they have the most reliable data as mentioned in Results. Beef has the best performance and veal is the worst (B): Box plot of feature ranks for the input features, where rank 1 is highest. Features are arranged in ascending order of average rank. Average ranks for both plant-based foods (and WH process) and animal-based foods (and DH process) are in the mid-range. No feature has a consistent high rank cross all the predictions. (C): Plots of performance-vs -#feature depict very different trends for prediction models, vitamin B6 and potassium are shown as examples for the WH process and vitamin B6 and zinc for the DH process. The best features for vitamin B6 (WH) are vitamin B6, zinc, vitamin C. Best Features for potassium (WH) are potassium, vitamin B9 (folate), water, magnesium, vitamin A, saturated fats, vitamin B5 (pantothenic acid), vitamin B1(thiamine), iron, poly unsaturated fats, selenium, vitamin B3 (niacin), vitamin B6 and zinc. Best Features for vitamin B6 (DH) are vitamin B6, magnesium, calcium, vitamin B2 (riboflavin), calcium, total fats, vit C and carbohydrates. Best features for zinc are zinc, phosphorus, calcium, potassium and total protein. The combined interpretation of B and C suggests that feature selection results differ for every nutrient prediction.

### High variability on the top predictive features

There is a notable lack of feature importance order across the prediction models. **Figure 3B** shows the feature ranks, where the features are ordered by their average rank across predictions. The average rank is in the mid-range for both the WH and DH process, suggesting that no feature has a consistent importance across all the predictions. **Figure 3C** shows performance by feature-size plots for vitamin B6 and potassium (WH) and vitamin B6 and zinc (DH) and the feature names are listed in the caption. The common observation is that the top ranked feature is the micronutrient itself in the raw food, as expected, but all other input features are specific to every prediction. The complete coverage of best features and feature ranks is in the **Supplementary materials**.

## DISCUSSION

In recent years the body of food composition data on preprocessing and postprocessing has grown, and so have simultaneously the expectations and vision [36] for big data in prediction tasks related to processing effects on food. But these expectations have yet to match the currently available computational capability and it is unknown whether the available data is sufficient for predictive capability and where the gaps might be. The results prove the potential of this existing data for predictive models and even show the categories that are more predictable namely leafy green vegetables in plant-based foods and beef in animal-based foods. A significant drawback in the data was the standard data convention of composition per 100grams of foods for raw and cooked foods which obscured the actual change and we remedied (but not reversed) this by the scaling method.

In the introduction we pointed out the discrepancies between the simpler prevalent methods and our ML models. Now, we elaborate on the discrepancies in perspective of our results, where the data scaling methods and ML model achieve an overwhelming improvement over the baseline. The scatter plots for vitamin B9(folate) and magnesium show that the baseline method overestimates the composition, which implies that the baseline RF is much greater than the RF inherent in the true data. RF represents the rate of loss which is influenced by process-related factors like processing times, surface area of vegetable exposed to processing conditions. Ideally for a fair comparison, these factors should be known for the baseline and matched to the data at hand. This can easily be addressed by recording additional meta-data. However, the more challenging discrepancy was that the baseline is a simple linear method, while the prediction model is a much more complex multiparametric non-linear ML model. Inevitably more sophisticated methods will emerge whether machine learning, mechanistic or a hybrid, and a suitable state-of-the-art baseline method will be available for comparison.

Our results show that methods of data analysis and predictive ML are valuable tools to assist in experimental design for food composition analysis, since the data generation process is time consuming and expensive. Specifically, we suggest how our methods justify decisions such as; the selection of food samples, recording of structured metadata/provenance, checking for data quality, and determining the composition feature set. The provenance of the data was incomplete in at least two different aspects. The composition data was calculated for some foods, and there was no explanation for the calculation method and no mention of the reference food /data used in the calculation method. It is unclear whether the samples for the raw and cooked food were related. Additionally, ontologies or structured vocabularies are a valuable resource when creating a format or structure for the dataset. Regarding data quality, we have described the anomalous condition in the **Results**. This is an example of a basic data sanity check, and especially in the context of a prediction hypotheses. Predictive performance depends on both the sample size as well as the entropy of the dataset, and one can use the predictive performance of the model as a guide for the sampling size for gathering new experimental data. There was only a single representative instance for each food and factors like geography, method of agriculture etc. are known to significantly impact the composition. Our results on prediction performance by category could justify the need for greater sampling. Regarding the feature space per sample, we suggest including process parameters and features known to influence nutrient loss such as pH.

Finally, representation of the composition per 100g of food, further obscured the data. We mitigated this issue by applying the data scaling methods, however our observation show that this is not a complete resolution and new standards for data representation are required. The results from applying the scaling methods on the composition data, has two unrelated interpretations; the effect on the size of non-anomalous food-pairs (**Figure 3**) and the effect on model performance trained on this data (**Supplementary materials**).

As seen in **Figure 3**, there is no significant effect (p-value =0.06) for the plant-based foods where the data representation causes a dilution bias, and the anomaly could instead be due to different food samples used for the raw and cooked analysis. Whereas there is a significant effect (p-value <0.01) on animal-based foods where the anomaly is due to a concentration bias. Regarding the prediction performance, a few additional components used in the PINS method had good results besides the hypotheses. For plant-based foods, the performance for SCS data was the best, followed by carbohydrate PINS data. For animal-based foods, the performance by PINS-proteins data was the better than for zinc, iron and cholesterol. However, the results for PINS-carbohydrate and PINS-protein are likely due to the methods used for generating this data. This analysis presents several questions for future inquiry and experimental validation, though the most important might be to ascertain a process-invariant nutrient and under which conditions and the biochemical/mechanistic explanation. This information might help for data transformations of existing data, but new data representation standards need to be considered and applied to future data generation efforts.

Food composition data is poised to grow in size and quality [37] [38] [39], and such predictive modelling applications can assist consumers in making reliable dietary decisions. For example, to choose food ingredients and cooking methods by getting answers to queries like-“How do boiling/streaming affect the nutrient profile of Vegetable X compared to roasting or frying”. Personalized health models must consider additional factors like bioavailability and individual physiological response, however dietary composition is an essential step towards that goal.

## AUTHOR CONTRIBUTIONS

T.N. gathered the data and performed all computational analysis. T.N. and I.T. wrote the manuscript. I.T. and T.N. designed the computational methods. I.T. conceived and supervised all aspects of the project.

## ACKNOWLEDGEMENTS

We would like to thank the members of the Tagkopoulos Lab, particularly Jason Youn and Gabriel Simmons for their feedback in machine learning methods, the USDA ARS methods and applications of food composition lab personnel and Dr. Bruce German for their suggestions. This work has been been supported by the USDA-NIFA AI Institute for Next Generation Food Systems (AIFS), USDA-NIFA award number 2020-67021-32855 and USDA grant 58-8040-8-015 to I.T.

